# *MiR-144* regulates adipogenesis by mediating formation of C/EBPα-FoxO1 protein complex

**DOI:** 10.1101/2021.11.11.468339

**Authors:** Weimin Lin, Xianyu Wen, Xuexin Li, Lei Chen, Wei Wei, Lifan Zhang, Jie Chen

## Abstract

Excessive adipogenesis caused obesity, which was a serious risk of health and led to a series of diseases, including type II diabetes (T2D) for example. Adipocyte as the basic unit of adipose tissue has emerged as one of significant target of the treatment of obesity-related metabolic syndromes by revealed its adipogenic molecular mechanism. MicroRNAs (miRNAs) have been demonstrated involving adipogenesis, and played a crucial role in the competitive endogenous RNA (ceRNA) effect. Besides that, C/EBPα as a crucial adipogenic regulator still lacked epigenetic explanation during pre-adipocyte adipogenesis. In this study, we first verified FoxO1 was one of the ceRNA of C/EBPα. They co-regulated adipogenesis through formed a protein complex that directly bound to its promoter to activate *AdipoQ*, and AdipoQ (Adiponectin) was a negative adipocytokines that suppressed adipogenesis, which played an important role in retaining adipogensis balance. Moreover, an adipose tissue specific enriched miRNA, miR-144 was the key regulator of the ceRNA effect between C/EBPα and FoxO1, which mediated the C/EBPα-FoxO1 complex formation, thus altered AdipoQ, furthermore regulated pre-adipocyte adipogenesis. This research will provide a new supplementary idea of the C/EBPα epigenetic role in pre-adipocyte adipogenesis.

## Introduction

Obesity threatens human health, which leads to a series of diseases, for instance, type II diabetes (T2D), hypertension, cardiovascular disease and different kinds of immunological diseases. In particular, by the report from Lancet, for children and adolescents aged from 5 to 19 years old, the rate of obesity nearly increased 40 times from 1975 to 2016 (1). Adipocyte as the basic unit of adipose tissue, its proliferation and differentiation lead to adipose tissue expansion, while excessive adipose tissue further results in obesity-related metabolic syndromes. In the clinical setting, adipocytes are emerging as a significant target of the treatment of obesity-related metabolic syndromes (2). The adipogenic molecular genetic mechanism has achieved great progress in past decades, helped us to understand the classic genetic mechanism, which involved peroxisome proliferator-activated receptor (PPARs) and CCAAT/enhancer binding proteins (C/EBPs) family (3, 4). In brief, in the early period of adipogenesis, dexamethasone (DEX) and isobutyl-methylxanthine (IBMX) promote C/EBPδ and C/EBPβ, then both of them *trans*-activate PPARγ and C/EBPα and further improve adipogenesis (5). PPARγ has been confirmed regulating insulin sensitivity (3, 6, 7). C/EBPα regulates the transcription of genes, whose products take part in programming the gluconeogenic pathway (8). Meanwhile, both of them are also involved in protein-complex formation, thus playing a regulatory role in biological progress regulation. For instance, the PPARγ-p53 complex has demonstrated activated pulmonary artery endothelial cell related genes transcription, thus contributed to a coordinated vasculoregenerative response (9). The PPARγ-PRDM16 complex in BAT that from HFD-fed *Ins1* ^+/-^ mice was increased, thus activated Sirt1 activity and increased Sirt3 abundance furthermore promoted UCP1 protein, which benefited adipose tissue metabolism by reprogramming its efficiency toward increased energy expenditure(10). The PPARγ-NcoR complex benefited vascular endothelium insulin resistance (11). Besides, C/EBPα-p50 complex has identified regulating *nfkb1* and displacing histone deacetylases NF-κB p50 homodimers thus regulating leukemia (12). The C/EBPα-FoxO1 complex regulates adiponectin expression, thus influences adipogenesis (13, 14). Both PPARs and C/EBPs families all belonged to the nuclear regulator that played transcriptional factor role among kinds of biological progress, the mechanism of complex still played their transcriptional role by binding to the promoter of their target genes. Moreover, by report, the abundance of single protein influenced the complex formation, which further identified the balance of protein abundance among protein complex was the key factor to regulate complex formation thus influence their transcriptional role for target genes (10).

MicroRNA (miRNAs), a short non-coding RNA with ~21 nucleotides, have been identified involving adipogenesis at the post-transcriptional regulation. Single miRNA targets hundreds of genes that share the same miRNAs response elements (MREs), which can complementary base pair with “seed sequence” within miRNAs. Hence, Pandolfi lab proposed a competitive endogenous RNA (ceRNA) hypothesis in 2011 (15, 16). From then on, multiple studies have proved the hypothesis.

As the report, there were a series of miRNA both targeted C/EBPα and FoxO1, meanwhile, C/EBPα-FoxO1 has been demonstrated forming the protein complex thus regulating adipogenesis by targeting AdipoQ promoter (13, 14, 17). On the other hand, the balance of abundance of single protein influenced complex formation, thus, in this research, we try to study the ceRNA effect whether regulated the interaction between *C/EBPα* and FoxO1, and to identify the key miRNA of ceRNA effect between C/EBPα and FoxO1, moreover to identify the regulatory role of C/EBPα-FoxO1 in adipogenesis.

## Materials and Methods

### Experiment animals

All animal experiments including the pre-adipocyte collection were approved and reviewed by the Animal Ethics Committee of Nanjing Agricultural University (STYK (Su) 2011-0036), and preformed according to the Regulations for Administration of Affairs Concerning Experimental Animals, China (11/14/1988). The Erhualian piglets all came from the Changzhou Erhualian Pig Production Cooperation (Changzhou, Jiangsu, China).

### Cell culture, transfection and differentiation

Isolating the subcutaneous adipose tissue from fresh slaughtered piglets, and soaking in phosphate-buffered saline (PBS), then transferring it to aseptic table to wash thrice with PBS (1% penicillin-streptomycin). Cutting up the adipose tissue then digesting them fully by collagenase type I (Invitrogen, Carlsbad, CA, USA) staying at 37℃ shaking bath (50rpm/min, more 2 hours). Adding equal volume of growth medium (89% Dulbecco’s modified Eagle’s medium/Ham’s F-12 (DMEM-F12), 10% fetal bovine serum (FBS) and 1% penicillin-streptomycin) to stop the digestion. Then, filtration through a 200μm nylon mesh for removing the undigested residue. The filtrated solution is dividing into 15ml centrifuge tubes, centrifuging them twice at 1,000 rpm/ min for 10 minutes to collect the pre-adipocyte. In the second time, abandoning the supernatant then adding 4ml growth medium to wash the pre-adipocyte, subsequently, culturing the pre-adipocyte in growth medium in cell incubator (37°C, 5% CO_2_). The medium was replaced every two or three days to ensure cell growth.

The cells are seeded into 6-well or 12-well plates for 24-48 hours until its density of 85% confluence. Then use the Lipofectamine 3000 (Invitrogen) following the protocol to transfect. All the vectors, siRNAs and miRNA mimics and inhibitors are shown in Supplementary Table S1.

To induce differentiation of porcine pre-adipocytes by using the adipogenic differentiation inducer medium (DIM) to stimulate for 8 days. The constituted of DIM are 2.5µM dexamethasone, 8.6µM insulin, 0.1mM 3-isobutyl-1 methylxanthine (IBMX), 1% penicillin-streptomycin and 10% FBS in Dulbecco’s modified Eagle’s medium/Ham’s-High Glucose (DMEM-HG) (Sigma-Aldrich, Shanghai, China). The medium is replacing it with a fresh one every two days.

### RNA isolation, library preparation, RT-qPCR

Isolating the porcine total RNA by Trizol reagent (TaKaRa, Dalian, China).

The mRNA and miRNA cDNA libraries are reverse-transcribed by PrimeScript™ RT Master Mix (TaKaRa, Dalian, China) and miRNA 1^st^ Strand cDNA Synthesis Kit (by stem-loop) (Vazyme, Nanjing, China). Using the AceQ Universal SYBR qPCR Master Mix (Vazyme, Nanjing, China) and the miRNA Universal SYBR qPCR Master Mix (Vazyme, Nanjing, China) to perform the RT-qPCR. The relative level of mRNA and miRNA expression estimate are 2^-ΔΔCt^ methods.

Every sample is performed in triplicate to test at least, while normalizing them to *GAPDH* and *miR-17* (18, 19), respectively. All the primers are shown at Supplementary Table S1.

### Oil-red-O staining and triglyceride assay

Briefly, differentiated porcine pre-adipocytes are gently washing thrice with fresh 1×PBS, fixed in 4% paraformaldehyde for 30 minutes, and then washing thrice with deionized water. Staining for 30 minutes with 60% saturated oil-red-O, subsequently (0.5% oil-red-O melts in pure isopropanol for 10 hours at 37 ℃, and filtering with a 0.22μM filter thrice, finally, diluting 3:2 in water before using.) (Sigma-Aldrich, Shanghai, China). Subsequently, washing thrice with PBS, the representative images of treating cells are captured by Zeiss Axiovert 40 CFL inverted microscope (Thornwood, NY, USA).

Total triglyceride of the pre-adipocyte is quantified by elution of oil-red-O with isopropanol and measuring the Absorbance at 510 nm wavelength.

### Western blotting

Lysing porcine pre-adipocyte and extracting the total protein by radioimmunoprecipitation assay (RIPA) lysis buffer (Vazyme, Nanjing, China) following the protocol. The protein is quantified by the BCA Protein Assay Kit (Beyotime Biotechnology, Jiangsu, China). Loading the protein onto the 12% SDS-PAGE, every protein sample is 15μg/well. After one-hour electrophoresis, the protein sample is transferred to the polyvinylidene difluoride (PVDF) membrane (Millipore, Billerica, USA). Blocking the membrane with 5% BSA (Bovine serum albumin), overnight incubation with primary antibody at 4℃, subsequently. After membrane washing the appropriate secondary antibody is used to incubate. The ECL Chemiluminescence Detection Kit (Thermo Scientific, USA) is used to detect and images are captured by the VersaDoc 4000 MP system (Bio-Rad, USA). The antibodies respectively are anti-C/EBPα (ZEN BIO, 383901), anti-FoxO1 (ZEN BIO, 383312), anti-AdipoQ (ZEN BIO, 383390).

### Bioinformatics analysis

Using the Targetscan (http://www.mirbase.org/) prediction tool to predict the *miR-144-3p* targeted genes. The sequence of precursor and mature *miR-144* are obtained from miRBase (http://www.mirbase.org/).

### Luciferase reporter assay

The wild and mutation sequence of purpose 3’UTR are shown at Supplementary TableS1. Transfecting the Luciferase reporter vectors contains purpose sequence into the pre-adipocyte, which are seeded in 12-well plates until it’s cultured at density of 85% confluence using the Lipofectamine 3000 (Invitrogen) following the protocol. Firefly and Renilla luciferase activity is used to quantify by Dual-Luciferase Reporter Assay System (Promega, Madison, WI, Germany).

### Co-immunofluorescence staining

For co-IP, the pre-adipocytes is washed thrice by PBS, then using the radioimmunoprecipitation assay (RIPA) lysis buffer (Vazyme, Nanjing, China) that contains 1% phenylmethylsulfonyl fluoride (PMSF) to lyse the cell, whole cell lysate is incubating with antibodies conjugated Protein A/G PLUS-Agarose (Santa Cruz, sc-2003) in RIPA at 4°C overnight. The antibodies respectively are anti-C/EBPα (ZEN BIO, 383901) and anti-FoxO1 (ZEN BIO, 383312). After the overnight incubation, washing the immunoprecipitates thrice with the fresh RIPA before subjection to immunoblotting (IB).

### Chromatin immunoprecipitation assay PCR (ChIP-PCR)

Using the ChIP Assay Kit (Boytime, Nanjing, China) following the manufacturer’s instruction to detect ChIP-PCR. In brief, using 1% formaldehyde to cross-link porcine pre-adipocyte at 37℃ for 10 minutes. 1×glycine is used to quench the cross-linked porcine pre-adipocyte at room temperature for 5 minutes. Using VCX750, the ultrasonic cell smash machine (Sonics, United States) to smash porcine pre-adipocyte and obtain DNA fragments. Subsequently, following the rule that per 100ml cell lysis buffer is incubating with 1 mg of anti-C/EBPα (ZEN BIO, 383901) and anti-FoxO1 (ZEN BIO, 383312) at 4℃ overnight, 60ml Protein A Agarose/SalmonSperm DNA is used to isolate immunoprecipitated complexes at 4℃ for 4 hours, washing the complexes as following: low salt wash buffer, high salt wash buffer, LiCl wash buffer once, and TE buffer twice. Final ChIP DNA fragments are subjected to PCR analysis using an *AdipoQ* promoter specific primers, as shown at Supplementary Table S1.

### Statistical analysis

SPSS software (21.0 version) is used to analyze the data. The data presents as mean ± standard error (s.e.m). Two-tailed Student’s t-test is used to assess the significance of the comparison between the groups, *P* < 0.05 is observed as a significant statistical difference.

## Results

### The ceRNA effect influenced C/EBPα-FoxO1 formation

C/EBPα and FoxO1 have been identified co-regulating gluconeogenesis and adipogenesis by forming the C/EBPα-FoxO1 protein complex (13, 14). Moreover, a series of miRNAs targeted C/EBPα and FoxO1 as well (2, 20–25). Thus, we first verified the ceRNA effect between C/EBPα and FoxO1 by detecting their mRNA expression pattern according to the ceRNA effect rule. First, we respectively transfected their 3’UTR or siRNAs into pre-adipocyte, then detected the expression patterns of *FoxO1* and *C/EBPα*. We found the *FoxO1* expression patterns were consistent with *C/EBPα* (Fig. 1A–D), *vice verse*. To be specific, take C/EBPα*-*3’UTR transfected for example, when we transfected C/EBPα-3’UTR into pre-adipocyte, C/EBPα-3’UTR sponged more abundant miRNA that targeted *FoxO1*, thus released their inhibition for FoxO1. On the contrary, when we knocked down *C/EBPα* by its siRNA, which led to miRNAs inhibition for *FoxO1* was up-regulated thus reduced *FoxO1* expression.

**Fig 1.**
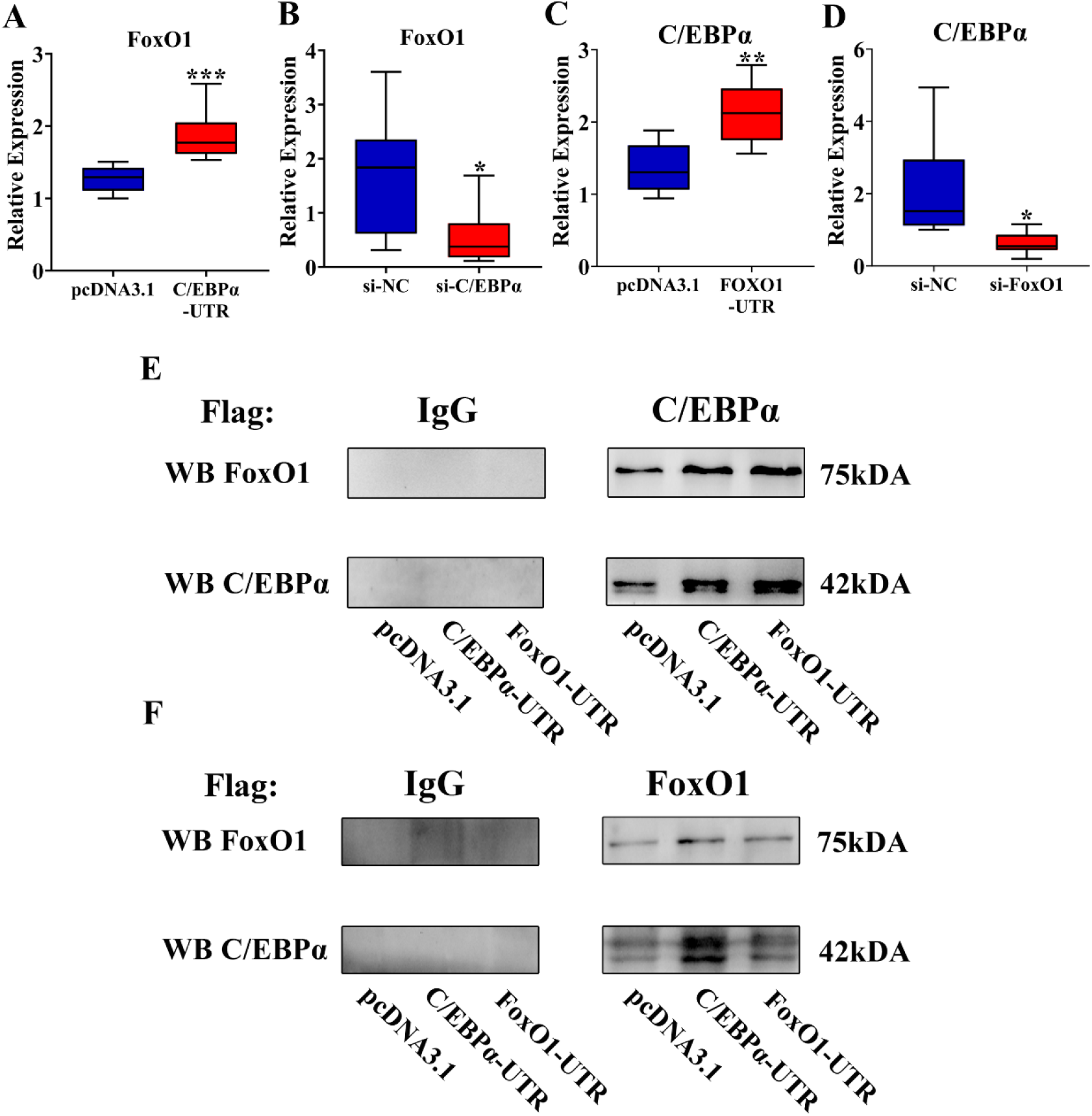
The ceRNA effect influenced C/EBPα-FoxO1 formation. (A-B) The FoxO1 relative expression after transfecting C/EBPα-3’UTR (A) or si-C/EBPα (B) into pre-adipocyte. (C-D) The C/EBPα relative expression after transfecting FoxO1-3’UTR (C) or si-FoxO1 (D) into pre-adipocyte. (E) IP analysis of interactions between C/EBPα and FoxO1 in pre-adipocyte after transfecting C/EBPα-3’UTR and FoxO1-3’UTR, used the anti-C/EBPα as the flag antibody, then immunoblotting (IB) analysis for the C/EBPα and FoxO1 protein level. (F) Used anti-FoxO1 as the flag antibody to detect the C/EBPα and FoxO1 protein level. **\*** means *P* < 0.05, **\*\*** means *P* < 0.01, **\*\*\*** means *P* < 0.001. Each bar indicates mean ± s.e.m.

Furthermore, we further identified whether the C/EBPα-FoxO1 protein complex level was regulated by the ceRNA effect. Hence, we used co-IP to detect their protein complex level. The result showed when we used the C/EBPα anti-body to pull-down the protein complex, the protein level of C/EBPα and FoxO1 was up-regulated in CEBPA-3′UTR and FoxO1-3′UTR groups compared with control group (Fig. 1E). Meanwhile, we used the FoxO1 anti-body to prove the results, which further demonstrated the complex protein level of both 3’UTR groups was increased compared with the control group, which was consistent with the result of C/EBPα anti-body (Fig. 1F).

### Identification of miRNA in the C/EBPα-FoxO1 ceRNA effect

We used the TargetScan prediction tool to predict the miRNAs binding sites within *C/EBPα*-3’UTR and *FoxO1*-3’UTR (Fig. 2A), meanwhile, the miRNAs usually have distinct tissue expression patterns (26, 27), we further identified the WAT specific one among the miRNAs that targeted C/EBPα and FoxO1. By RT-qPCR detection, we found the *miR-144* was the highly expressed miRNA in WAT by comparing with other miRNAs, showed in Fig. 2B. However, *miR-144* has been elucidated targeting human and murine *C/EBPα* and *FoxO1* gene, however, whether the *ssc-miR-144* targeted porcine *C/EBPα* and *FoxO1* still need to be further identified. Thus, we constructed the pmirGLO dual-luciferase reporter vector respectively containing *miR-144* binding sites of *C/EBPα* and *FoxO1* 3’UTR, which included 300bp up- and downstream sequence of binding sites. Dual-luciferase reporter assay system results illustrated *ssc-miR-144* targeted porcine C/EBPα and FoxO1 (Fig. 2C–D). We respectively detected relative expression level of *C/EBPα* and *FoxO1* gene after either transfecting *ssc-miR-144* mimics or inhibitors into pre-adipocyte, the results identified *ssc-miR-144* inhibited C/EBPα and FoxO1(Fig. 2E–F), and the western blotting results further proved that (Fig. 2G). To sum up, we found that *ssc-miR-144* targeted C/EBPα and FoxO1 thus suppressed their mRNA and protein expression.

**Fig 2.**
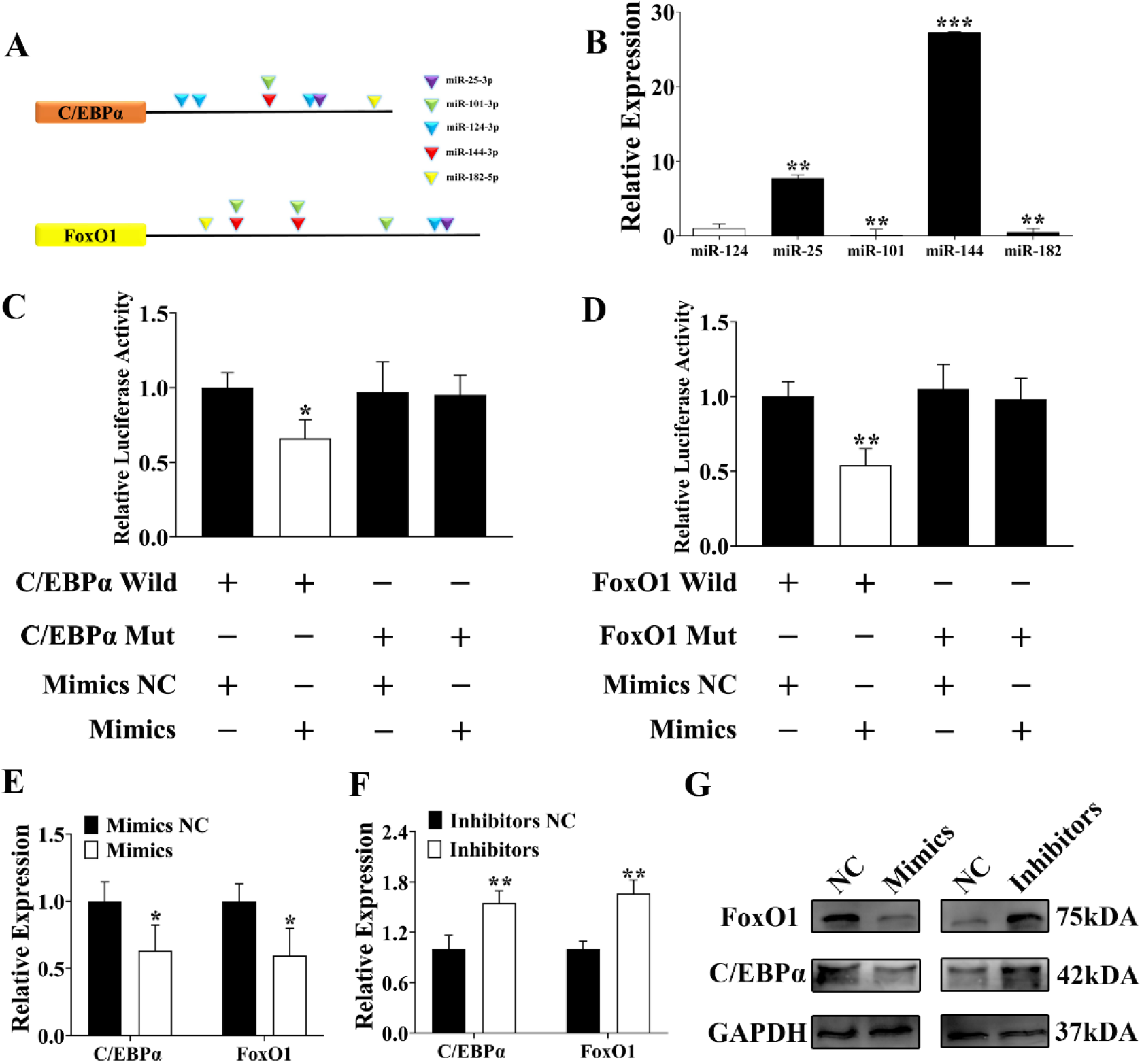
Identification of key miRNA in the C/EBPα-FoxO1 ceRNA effect. (A) The prediction binding sits of miRNAs within C/EBPα-3′UTR and FoxO1-3′UTR by TargetScan prediction tool. (B) The miRNAs relative expression within the WAT. The *miR-124* was regarded as the standard miRNA. (C-D) The relative luciferase activity of pmirGLO-C/EBPα-3’UTR (C) and pmir-GLO-FoxO1-3’UTR (D), which respectively contained wild or mutated binding site and co-transfected with miR-144 into pre-adipocyte. (E-F) The *C/EBPα* and *FoxO1* relative expression in pre-adipocyte after transfecting miR-144 mimics (E) or inhibitors (F). (G) The C/EBPα and FoxO1 protein level in pre-adipocyte after respectively transfecting miR-144 mimics or inhibitors. **\*** means *P* < 0.05, **\*\*** means *P* < 0.01, **\*\*\*** means *P* < 0.001. Each bar indicates mean ± s.e.m.

### *MiR-144* was the key regulator of the C/EBPα-FoxO1 ceRNA effect

The key mechanism of the ceRNA effect was the random and free sponging of miRNA by target genes (15, 16). As the Fig. 1 and Fig. 2 results showed *miR-144* was the WAT specific enriched miRNA meanwhile targeted C/EBPα and FoxO1 as well, thus, we further verified whether *miR-144* was the key regulator of the C/EBPα-FoxO1 ceRNA effect. We respectively co-transfected pcDNA3.1-C/EBPα-3’UTR or si-C/EBPα with *miR-144* inhibitors into pre-adipocyte. By RT-qPCR detection, we found the expression patterns of FoxO1 were weakened by *miR-144* inhibitors (Fig. 3A–B). Moreover, we also co-transfected pcDNA3.1-FoxO1-3’UTR or si-FoxO1 with *miR-144* inhibitors into pre-adipocyte, result showed C/EBPα expression patterns were weakened by *miR-144* inhibitors as well (Fig. 3C–D).

**Fig 3.**
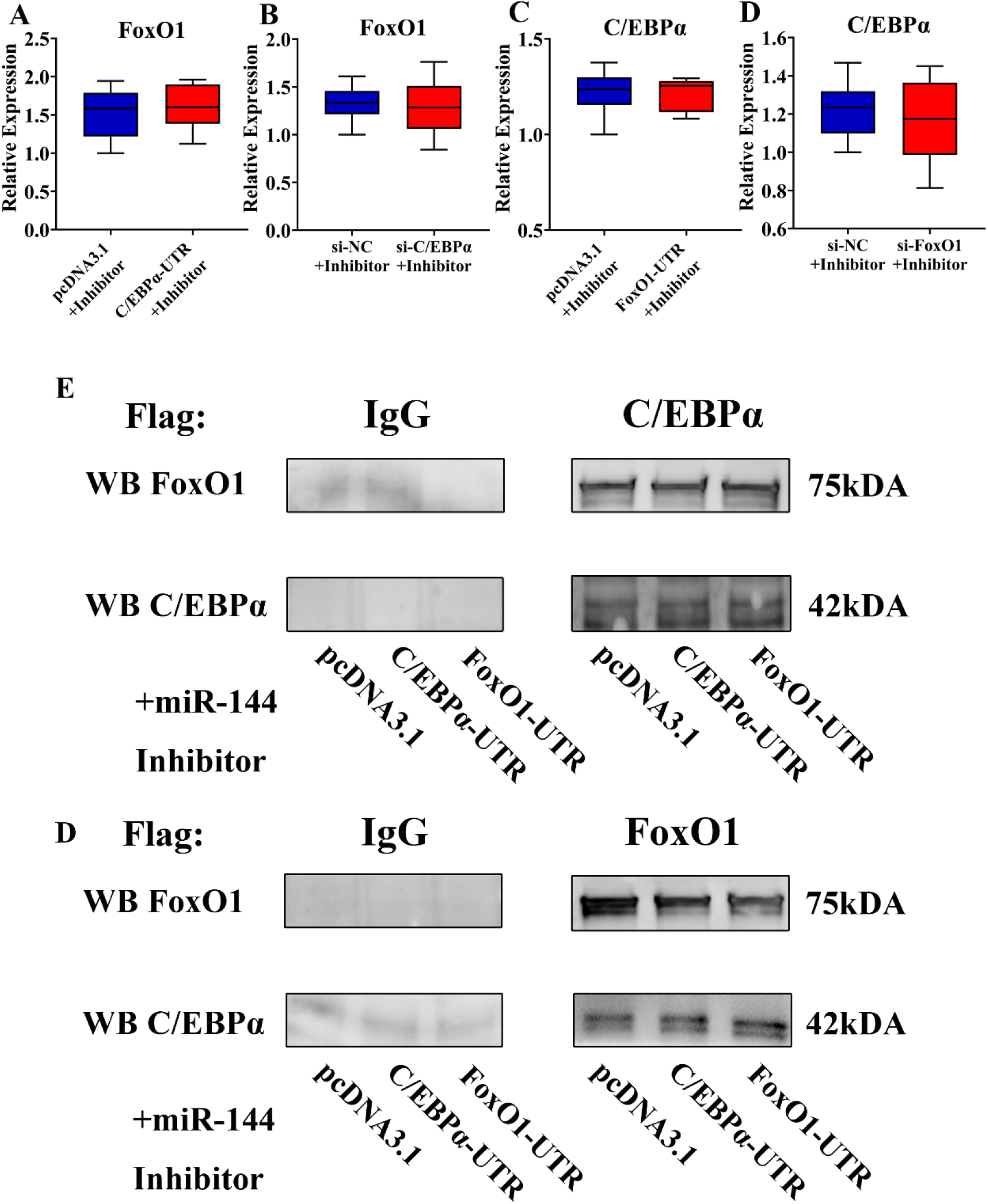
*MiR-144* was the key regulator of the C/EBPα-FoxO1 ceRNA effect. (A-B) The FoxO1 relative expression after co-transfecting C/EBPα-3’UTR (A) or si-C/EBPα (B) with miR-144 inhibitors into pre-adipocyte. (C-D) The C/EBPα relative expression after co-transfecting FoxO1-3’UTR (C) or si-FoxO1 (D) with miR-144 inhibitors into pre-adipocyte. (E) IP analysis of interactions between C/EBPα and FoxO1 in pre-adipocyte after co-transfecting C/EBPα −3’UTR and FoxO1-3’UTR with miR-144 inhibitors, used anti-C/EBPα as the flag antibody, then immunoblotting (IB) analysis of the protein level of C/EBPα and FoxO1. (F) Used the anti-FoxO1 as the flag antibody to detect the protein level of C/EBPα and FoxO1.

Besides, we also verified the C/EBPα-FoxO1 level after co-transfecting 3’UTR and *miR-144* inhibitors using co-IP. When we used anti-C/EBPα to pull down the C/EBPα-FoxO1 complex, we found the complex level was weakened by *miR-144* inhibitor (Fig.3E), on the other hand, when we used the anti-FoxO1 to further identify the C/EBPα-FoxO1 complex level, the result proved *miR-144* inhibitor eliminated the ceRNA effect of C/EBPα-FoxO1, thus the C/EBPα-FoxO1 complex level would not altered by 3’UTR transfection (Fig. 3F).

To sum up, we found that *miR-144* was the key regulator of C/EBPα-FoxO1 ceRNA effect, thus altering C/EBPα-FoxO1 complex formation.

### The ceRNA effect altered the C/EBPα-FoxO1 complex thus regulated AdipoQ

Adiponectin (AdipoQ) was secreted by mature adipose tissue in mammal, meanwhile, it was a negative adipocytokine that was involved in adipogenesis (17). C/EBPα and FoxO1 have been identified to co-regulate mice adiponectin expression by C/EBPα-FoxO1 protein complex (13). We have reported that the C/EBPα-FoxO1 protein complex bound to porcine *AdipoQ* promoter (17).

To explore whether the ceRNA effect of C/EBPα-FoxO1 altered the AdipoQ, we used the ChIP-PCR to identify the *AdipoQ* promoter that C/EBPα-FoxO1 complex bound to. The result indicated the 3’UTR transfected groups had more abundant pull-down promoter fragments compared with the control group (Fig. 4A). To be specific, in C/EBPα antibody pull-down result, we found the complex bound to more abundant *AdipoQ* promoter fragments in FoxO1-3’UTR transfected group. So did the C/EBPα-3’UTR transfected group in the FoxO1 antibody pull-down result (Fig. 4A–B). Besides, we further transfected the siRNAs of C/EBPα and FoxO1, results showed that both siRNAs groups reduced the pull-down promoter fragments than the control group, which due to the inhibition by siRNAs. However, in terms of the specific results, both ChIP-PCR results were still consistent with the ceRNA effect (Fig. 4C–D). We further detected the AdipoQ protein by transfecting 3’UTR into pre-adipocyte, we found the AdipoQ protein of C/EBPα-3’UTR and FoxO1-3’UTR groups were up-regulated compared with control group (Fig. 4E). Meanwhile, when we transfected siRNAs of C/EBPα or FoxO1 into pre-adipocyte, their AdipoQ protein were down-regulated compared with control group (Fig. 4E). These results indicated that the ceRNA effect influenced C/EBPα-FoxO1 protein complex formation, thus regulated the AdipoQ by binding to its promoter.

**Fig 4.**
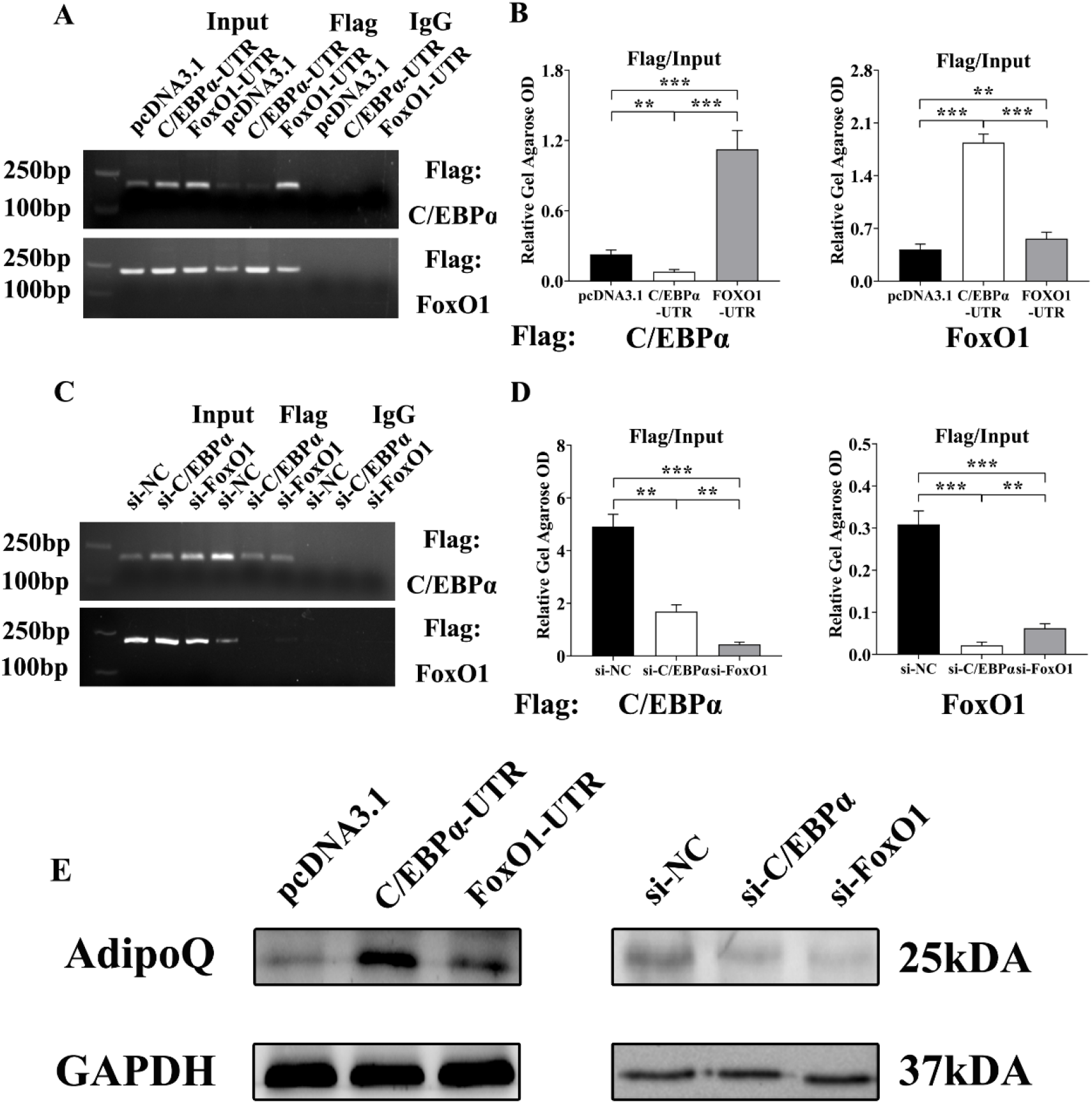
The ceRNA effect altered the C/EBPα-FoxO1 complex thus regulated AdipoQ. (A-B) ChIP-PCR detection of the C/EBPα-FoxO1 complex pull-down DNA fragments within the *AdipoQ* promoter region after transfecting 3’UTR into pre-adipocyte, respectively used the anti-C/EBPα and anti-FoxO1 as the flag antibody (A), and the OD value was visualized by the ImageJ software (B). (C-D) C/EBPα-FoxO1 complex pull-down promoter fragments in the siRNAs transfected pre-adipocytes (C), and the OD value (D). (E) The western blotting result of AdipoQ after respectively transfecting 3’UTR of C/EBPα and FoxO1 or their siRNAs into pre-adipocyte. **\*\*** means *P* < 0.01, and **\*\*\*** means *P* < 0.001. Each bar indicates mean ± s.e.m.

### MiR-144 played a crucial role in regulating AdipoQ

As the Fig. 4 identified, the C/EBPα-FoxO1 complex regulated AdipoQ by binding to its promoter, meanwhile, the ceRNA effect of C/EBPα-FoxO1 altered AdipoQ expression. Moreover, as the Fig. 2 showed, we found the *miR-144* was the crucial regulator for C/EBPα-FoxO1 complex formation. Hence, we further explored whether the *miR-144* altered AdipoQ by regulating the C/EBPα-FoxO1 ceRNA effect. We respectively co-transfected C/EBPα-3’UTR and FoxO1-3’UTR with *miR-144* inhibitors into pre-adipocyte. ChIP-PCR still was used to identify the *AdipoQ* promoter fragment that the C/EBPα-FoxO1 complex bound to. We found the *AdipoQ* promoter fragment of C/EBPα-3’UTR and FoxO1-3’UTR was weakened by *miR-144* inhibitors (Fig. 5A–B). Besides, we also co-transfected siRNAs of C/EBPα and FoxO1 with *miR-144* inhibitors into pre-adipocyte, then detected *AdipoQ* promoter fragments. The results showed the *AdipoQ* fragment that C/EBPα-FoxO1 complex bound to were recovered by *miR-144* inhibitors (Fig. 5C–D). Furthermore, we further detected the AdipoQ protein after co-transfecting 3’UTR or siRNAs with *miR-144* inhibitors. We didn’t find the AdipoQ protein level were altered compared with control groups (Fig. 5E). However, compared with Fig. 4, we found the alteration of AdipoQ protein level was eliminated by miR-144 inhibitor, in other words, miR-144 played a crucial role in regulating AdipoQ by ceRNA effect of C/EBPα-FoxO1.

**Fig 5.**
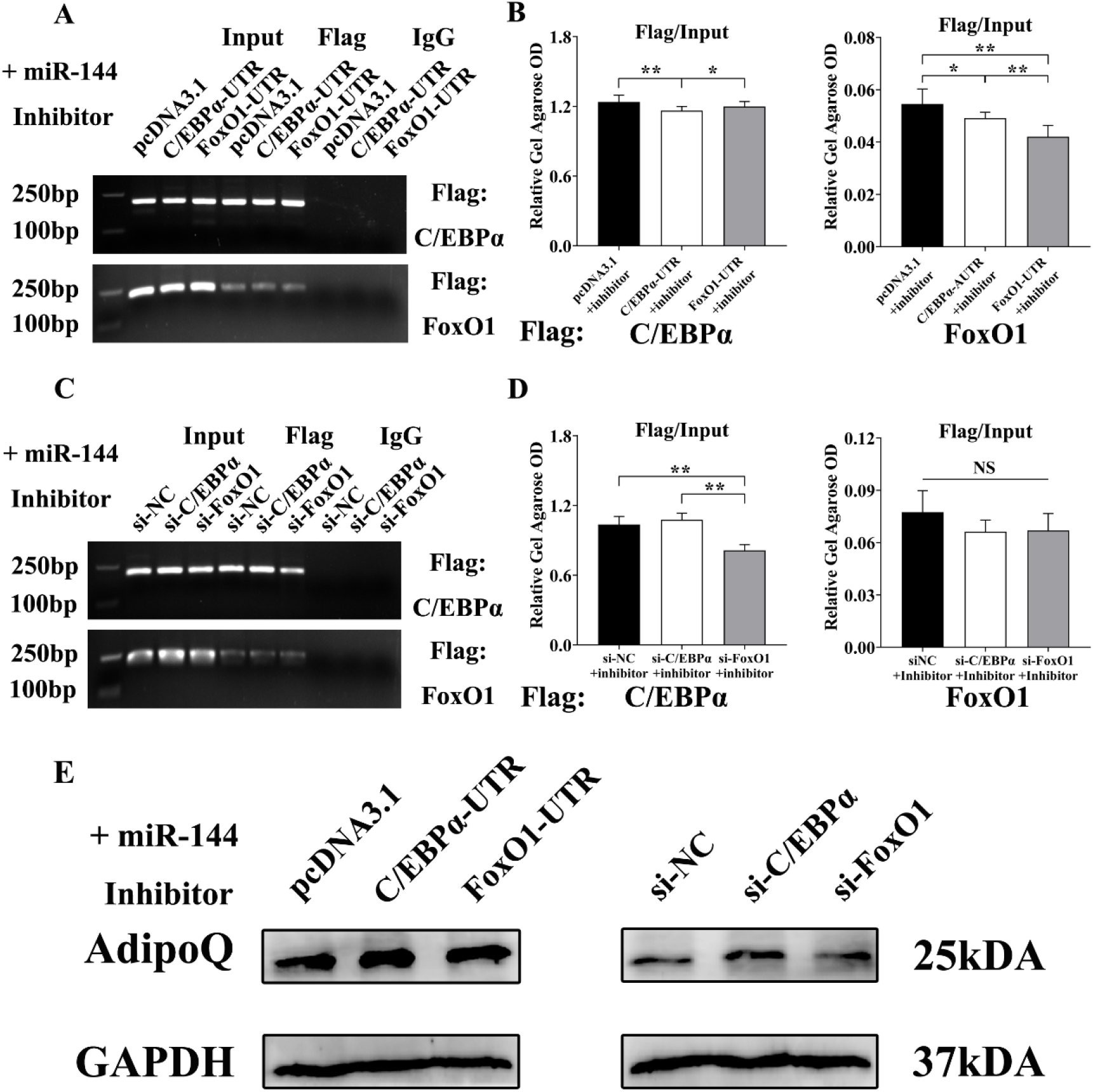
*MiR-144* played a crucial role in regulating AdipoQ. (A-B) ChIP-PCR detection of the C/EBPα-FoxO1 complex pull-down DNA fragments within the *AdipoQ* promoter region after co-transfecting 3’UTR with miR-144 inhibitors into pre-adipocyte, then respectively used the anti-C/EBPα and anti-FoxO1 as the flag antibody (A), and the OD value was visualized by the ImageJ software (B). (C-D) ChIP-PCR detection of siRNAs and miR-144 inhibitors co-transfected pre-adipocyte (C), and the OD value (D). (E) The AdipoQ protein in pre-adipocyte after co-transfecting miR-144 inhibitors with 3’UTR or siRNAs. **\*** means *P* < 0.05, **\*\*** means *P* < 0.01. Each bar indicates mean ± s.e.m.

### The ceRNA effect altered AdipoQ thus regulated pre-adipocyte adipogenesis

As a negative adipocytokine, adiponectin has been identified suppressing adipogenesis, and the C/EBPα-FoxO1 complex regulated AdipoQ. Hence, we further induced pre-adipocyte differentiation for 8 days to figure out the adipogenic role of the ceRNA effect of C/EBPα-FoxO1. We found that the adipogenesis of pre-adipocyte in every group stayed at a low level on day 2, and was significantly up-regulated on day 8 (Fig. 6A–B). Along with the differentiated period of pre-adipocyte, the adipogenesis of pre-adipocyte in FoxO1-3’UTR group and C/EBPα-3’UTR group were suppressed compared with the control group, among them, the adipogenesis of C/EBPα-3’UTR was stayed at the lowest level (Fig. 6A–B). To further confirm adipogenesis of pre-adipocyte was regulated by the C/EBPα and FoxO1 ceRNA effect, as Fig. 5 results showed their ceRNA effect regulated AdipoQ, especially in C/EBPα-3’UTR transfected group (Fig.5A), thus we respectively detected the *AdipoQ* relative expression on day 2 and day 8. The results showed that AdipoQ was up-regulated in both 3’UTR transfected groups compared with control group on day 2 (Fig. 6C), especially, the *AdipoQ* of C/EBPα-UTR group was the highest, which was consistent with the Figure. 5 result, and it further demonstrated the ceRNA effect of C/EBPα-FoxO1 regulated adipogenesis. While, on day 8, we found the *AdipoQ* of every group stayed at the same level. By our consideration, probably for two reasons. First, it was the negative feedback regulation of mature adipocytes in late adipogenesis, the mature adipocyte was able to secrete adiponectin after all. Besides, another possible reason was the 8-day differentiated period was longer than the half-life period of the over-expressed 3’UTR sequence. Hence, we cannot detect the *AdipoQ* expression level regulated by the ceRNA effect.

**Fig 6.**
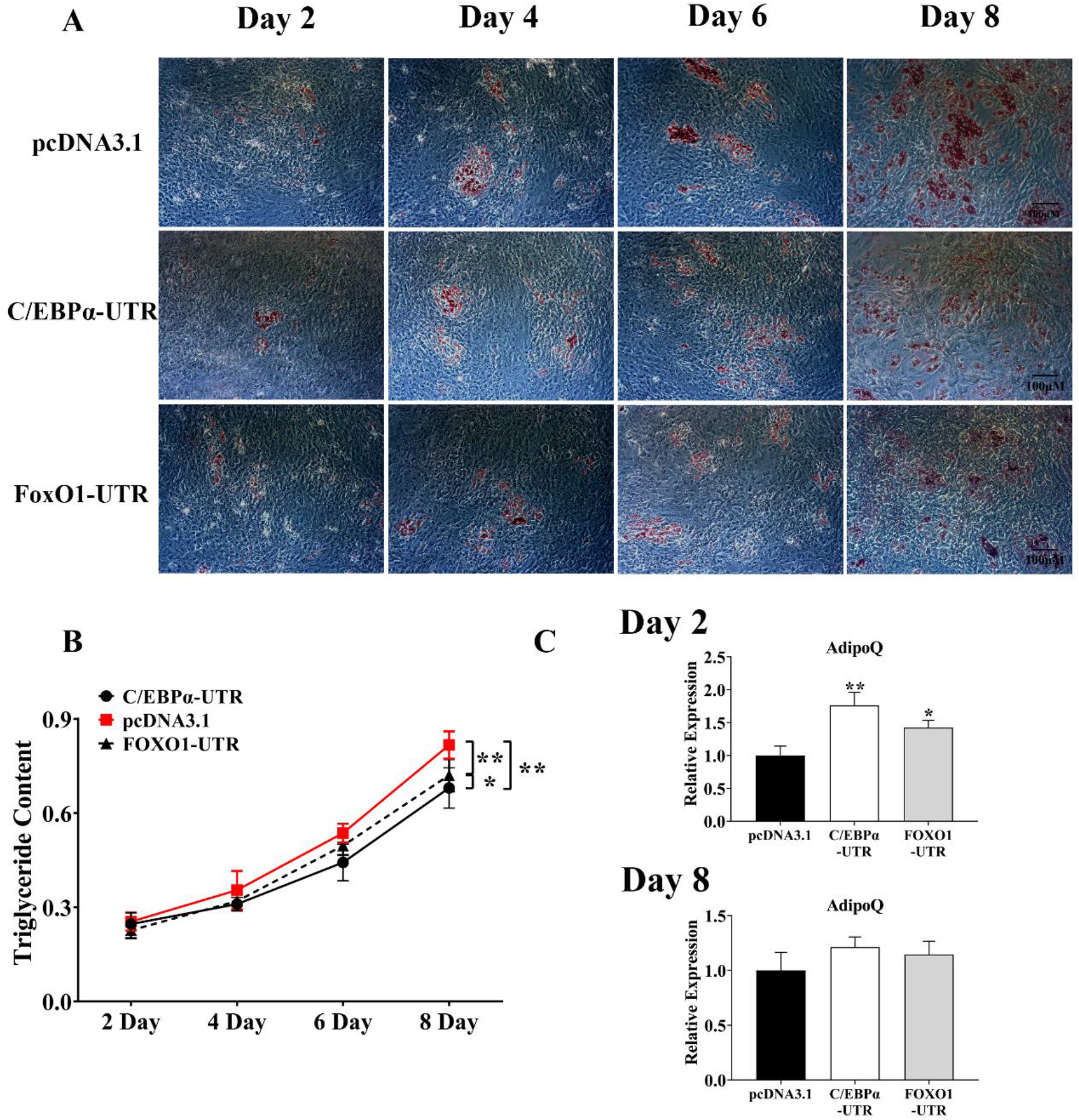
The ceRNA effect altered AdipoQ thus regulated pre-adipocyte adipogenesis. (A-B) The oil red O staining of 3’UTR transfected pre-adipocytes from day 2 to 8 and their triglyceride assay. Scale bars, 100μm. (C) AdipoQ relative expression in 3’UTR transfected pre-adipocytes, respectively detected on day 2 and day 8. **\*** means *P* < 0.05, **\*\*** means *P* < 0.01. Each bar and line indicates mean ± s.e.m.

### *MiR-144* was the crucial regulator of the ceRNA effect thus influenced adipogenesis

The Fig. 5 results showed that the ceRNA effect among *C/EBPα* and *FoxO1* regulated pre-adipocyte adipogenesis, and *miR-144* was the key component of the *C/EBPα*-*FoxO1* ceRNA effect. Hence, we co-transfected 3’UTR and *miR-144* inhibitors into pre-adipocyte, and detected the adipogenesis status from day 2 to day 8. We found that, even on day 8, the adipogenesis status of pre-adipocyte in every group all stayed at the same level, in brief, the ceRNA effect of 3’UTR that regulating pre-adipocyte adipogenesis was weakened by *miR-144* (Fig.7A–B). Which further proved that *miR-144* was key regulator for the *C/EBPα*-*FoxO1* ceRNA effect during pre-adipocyte adipogenesis. And we also detected the expressions of *AdipoQ* on day 2 and day 8, respectively. The results showed they were weakened by *miR-144* (Fig. 7C). The above results further proved the ceRNA effect of C/EBPα and FoxO1 was involved in adipogenesis, and the *miR-144* was the crucial regulator of their ceRNA effect thus regulated adipogenesis.

**Fig.7.**
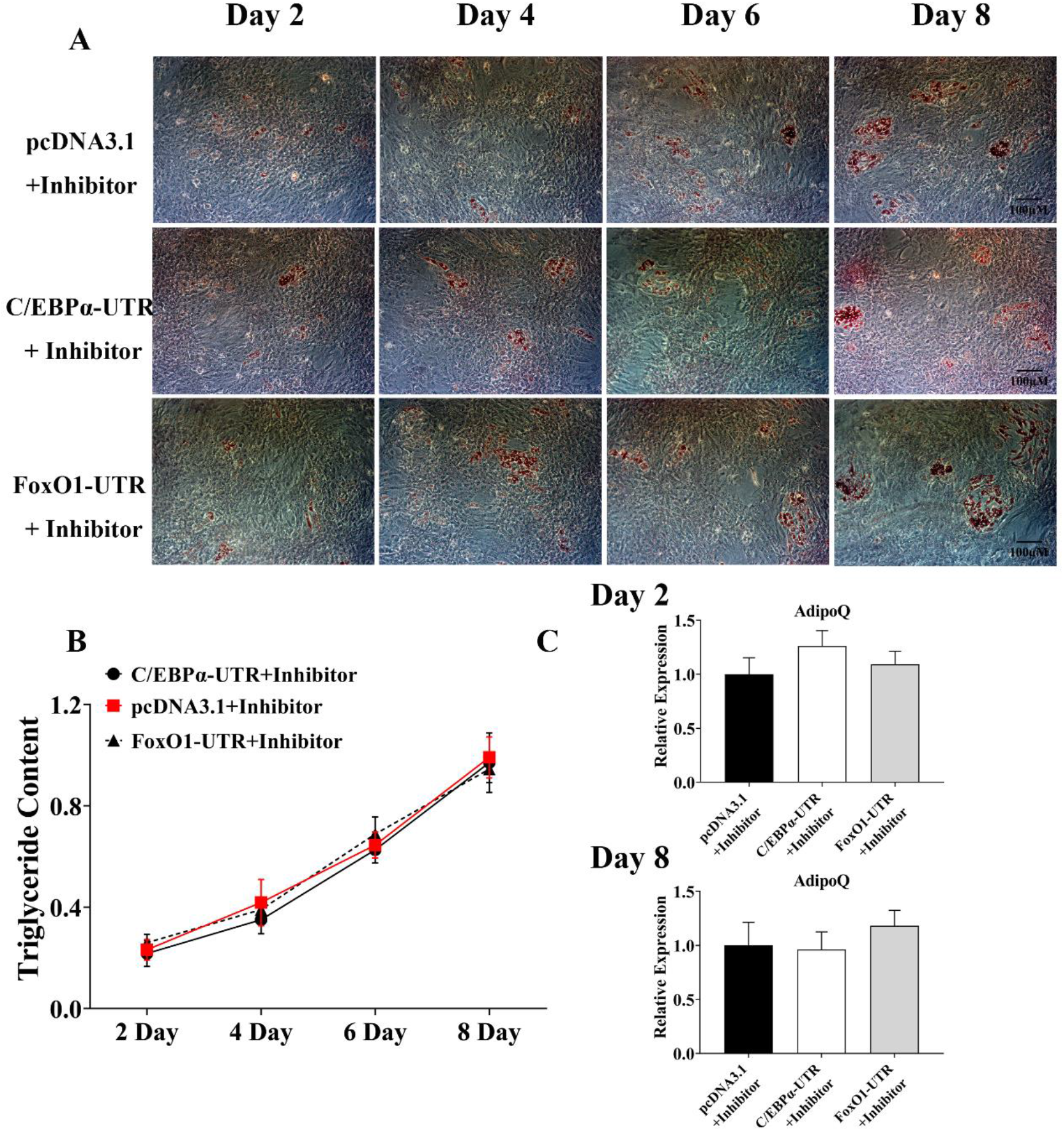
*MiR-144* was the crucial regulator of the ceRNA effect thus influenced adipogenesis. (A-B) The oil red O staining of 3’UTR and *miR-144* inhibitors co-transfected pre-adipocytes from day 2 to 8 and their triglyceride assay. Scale bars, 100μm. (C) *AdipoQ* relative expression of co-ransfected pre-adipocytes, respectively detected on day 2 and day 8. Each bar and line indicates mean ± s.e.m.

## Discussion

Obesity is the source of most metabolic diseases, including T2D for example. To make matters worse, the obesity rate of children and adolescents becomes more and more serious year by year. By a report, approximately 40% of the population of the world suffers from the risk of obesity, among them, nearly 20% are adolescents (28). Hence, the exploration of adipogenic molecular mechanisms is necessary and urgent for clinical treatment. As the basic unit of adipose tissue, adipocyte is regarded as a potential target for obesity-related metabolic syndromes. Generally, C/EBPα and PPARγ are considered to be the master regulators of adipogenesis. PPARγ is involved in both white and brown adipocyte formation. Whereas, C/EBPα still lacks evidence of regulating brown adipocytes development (29, 30). Moreover, both PPARs and C/EBPs families belonged to the nuclear regulator, which also have been reported involving in protein-complex formation, thus playing a regulatory role in a series of biological progresses, including adipogenesis. Among them, the C/EBPα-FoxO1 complex has been demonstrated binding to AdipoQ thus playing a transcriptional role (13, 14). Besides, the balance of protein abundance among protein complex was the key factor to regulate complex formation, hence we further verified whether single protein was regulated by the protein abundance of another one thus altered the complex formation. As the reports, kinds of miRNAs were shared between C/EBPα and FoxO1, based on the ceRNA effect, we respectively transfected their 3’UTR into pre-adipocyte, then detected the C/EBPα-FoxO1 complex level. We found that the complex level was altered by the 3’UTR transfection (Fig 1). For detail, C/EBPα-3’UTR promoted the FoxO1 protein level, thus increasing their complex level. While FoxO1-3’UTR up-regulated the C/EBPα protein level, thus improved the complex level. The results one side identified the abundance of both proteins determined their complex level, another side which demonstrated C/EBPα and FoxO1 were acting as the ceRNA of each other. Based on the rule of ceRNA effect, miRNAs shared among their targets determined whether the ceRNA effect works or not (15, 16). According to our detection, *miR-144*, a WAT specifically expressed miRNA, targeted C/EBPα and FoxO1 (Fig. 2). Thereby, we further explored the role of *miR-144* among C/EBPα-FoxO1 ceRNA effect. We co-transfected its inhibitors and both 3’UTR into pre-adipocyte, result showed the alteration of the C/EBPα-FoxO1 complex was recovered (Fig. 3). The ceRNA effect showed when C/EBPα-3’UTR sponged more abundance miRNAs, thus released their inhibition of for FoxO1, which further promoted FoxO1 expression, vice versa. As our results showed miR-144 inhibitor destroyed the ceRNA effect of C/EBPα-FoxO1. Which proved miR-144 was the crucial regulator that regulated C/EBPα-FoxO1 complex formation.

Furthermore, the C/EBPα-FoxO1 complex has been demonstrated regulating AdipoQ by binding to its promoter (13, 14). Adiponectin as a negative adipocytokine, was synthesized and secreted by the mature adipose tissue, which benefited to keep the endogenous adipogenic balance of adipogenesis in vivo (31). Hence, whether the ceRNA effect of C/EBPα-FoxO1 was involved in AdipoQ regulation, we further verified that. The results showed the AdipoQ protein was up-regulated in both 3’UTR transfection groups compared with the control group. In particular, the FoxO1 was the main functional regulator among C/EBPα-FoxO1 complex that targeted AdipoQ (13). Hence, the AdipoQ protein level of si-FoxO1 group should be the lowest compared with the other groups. However, we found the AdipoQ protein level of si-FoxO1 group was no alteration compared with other groups. By the Fig.2 showed, the miR-144 targeted FoxO1, when we co-transfected miR-144 inhibitors and si-FoxO1 into pre-adipocyte, the inhibition effect of miR-144 for FoxO1 was weakened by miR-144 inhibitors, thus led to the FoxO1 protein level was not alteration compared with control group (Fig. 5E). Furthermore, the results of the Fig.4 to Fig.5 further identified that the miR-144 was the crucial regulator that determined whether the ceRNA effect works or not, thus regulated AdipoQ (Fig. 4–5).

We further verified whether the ceRNA effect regulated pre-adipocyte adipogenesis. We transfected both 3’UTR into pre-adipocyte as well, after 8-day differentiation induced, the adipogenesis of every group was detected, we found the adipogenesis of both 3’UTR were down-regulated compared with the control group, which due to their promoting effect on AdipoQ thus inhibited adipogenesis (Fig. 6). However, to our surprise, we only found the promoting effect of 3’UTR on AdipoQ on day-2 rather than day-8. We hold the opinion that there maybe two probable reasons. First, the half-life of 3’UTR vector was shorter than the induced differentiation period (32, 33), which caused the ceRNA effect was weakened in day-8. Besides, the other reason perhaps the feedback of the mature adipocytes, they synthesized and secreted the adiponectin thus weakened the ceRNA effect promotion for AdipoQ. After all, the AdipoQ expression pattern related with the adipogenesis, which was up-regulated with the increase of the differentiation period (34, 35). Meanwhile, we co-transfected the 3’UTR and *miR-144* inhibitor into pre-adipocyte then induced the pre-adipocyte adipogenesis. Our results showed the ceRNA effect of C/EBPα-FoxO1 was destroyed by *miR-144* inhibitor thus led to their promotion for AdipoQ was weakened as well, furthermore caused the adipogenesis of both 3’UTR group were recovered (Fig. 7).

## Conclusion

In summary, we verified C/EBPα-FoxO1 protein complex was altered by their ceRNA effect, and the *miR-144* was the crucial regulator of the ceRNA effect, furthermore, the complex induced *AdipoQ* by binding to its promoter, thereby regulated pre-adipocyte adipogenesis.

## Author contributions

Weimin Lin conceived the ideas, performed the analyses and wrote the manuscript; Weimin Lin and Xianyu Wen and Xuexin Li respectively implemented the package; Lei Chen contributed to data analysis and manuscript checked; Wei Wei and Lifan Zhang contributed to fully manuscript checked; Jie Chen conceived the ideas, supervised the project analyzed and contributed to manuscript preparation.

## Supplemental data

This paper contains Supplemental data on URL:https://figshare.com/articles/figure/Supplementary_Table_S1_docx/16817347 DOI: https://10.6084/m9.figshare.16817347

